# Invasive lobular carcinoma integrated multi-omics analysis reveals silencing of Arginosuccinate synthase and upregulation of nucleotide biosynthesis in tamoxifen resistance

**DOI:** 10.1101/2025.01.16.633236

**Authors:** Annapurna Gupta, Fouad Choueiry, Jesse Reardon, Nikhil Pramod, Anagh Kulkarni, Eswar Shankar, Steven T. Sizemore, Daniel G. Stover, Jiangjiang Zhu, Bhuvaneswari Ramaswamy, Sarmila Majumder

## Abstract

Invasive Lobular Carcinoma (ILC), a distinct subtype of breast cancer is hallmarked by E-Cadherin loss, slow proliferation, and strong hormone receptor positivity. ILC faces significant challenges in clinical management due to advanced stage at diagnosis, late recurrence, and development of resistance to endocrine therapy - a cornerstone of ILC treatment. To elucidate the mechanisms underlying endocrine resistance in ILC, ILC cell lines (MDA-MB-134-VI, SUM44PE) were generated to be resistant to tamoxifen, a selective estrogen receptor modulator. The tamoxifen-resistant (TAMR) cells exhibit a 2-fold increase tamoxifen IC_50_ relative to parental cells. Metabolomics and RNA-sequencing revealed deregulation of alanine, aspartate, and glutamate metabolism, purine metabolism, and arginine and proline metabolism in TAMR cells. Among the fifteen commonly dysregulated genes in these pathways, low *ASS1* expression was identified in the TAMR cells and was significantly correlated with poor outcome in ILC patients, specifically in the context of endocrine therapy. Our study reveals methylation mediated silencing of *ASS1* in TAMR cells as a likely mechanism of downregulation. Demethylation restored *ASS1* expression and correspondingly reduced tamoxifen IC_50_ toward parental levels. Nucleic acid biosynthesis is augmented in TAMR cells, evidenced by increase in nucleotide intermediates. Both TAMR cell lines demonstrated increased expression of several nucleic acid biosynthesis enzymes, including *PAICS, PRPS1, ADSS2, CAD, and DHODH*. Furthermore, CAD, the key multifunctional protein of *de novo* pyrimidine biosynthesis pathway is differentially activated in TAMR cells. Treating TAMR cell with Decitabine, a demethylating agent, or Farudodstat, a pyrimidine biosynthesis inhibitor, markedly augmented efficacy of tamoxifen. Collectively, our study unveils *ASS1* downregulation as a novel mechanism underlying acquired tamoxifen resistance in ILC and establishes a metabolic link between ASS1 and nucleic acid biosynthesis. Restoring *ASS1* expression or inhibiting pyrimidine biosynthesis restored tamoxifen sensitivity. *ASS1* could be a potential biomarker and therapeutic target in tamoxifen resistant ILC patients, warranting further investigation.

## Introduction

Invasive lobular cancer (ILC) is a distinct histological and molecular subtype accounting for ∼15% of all breast cancers and is the second most common subtype of invasive breast cancer after invasive ductal carcinoma (IDC) [1, 2]. Typically, ILC is estrogen receptor (ER) positive, progesterone receptor (PR) positive, and HER-2 negative with a low mitotic index. Loss of E-cadherin is a hallmark of ILC which contributes to the unique morphology, frequent multifocality, and characteristic metastatic pattern to serosa including ovaries, gut, and peritoneum [3, 4, 5]. Patients with ILC face delayed and higher stage at diagnosis, and long-term worse disease-free and overall survival [6, 7]. While endocrine therapy remains the cornerstone in treatment of ILC, one of the greatest hurdles in the management of ILC is resistance to endocrine therapy leading to late recurrences [8]. Though a prevalent subtype, ILC remains relatively understudied, and is frequently grouped with ER/PR positive IDC; consequently, ILC management including screening, treatment, and follow-up strategies are largely based on data from IDC. Hence there is an unmet need to address mechanism underlying endocrine resistance in ILC and develop strategies to overcome resistance.

Tamoxifen, a selective estrogen receptor modulator (SERM), is a cornerstone in endocrine therapy for premenopausal (and certain postmenopausal) women with estrogen receptor-positive (ER+) breast cancer [9]. There is active clinical development of novel SERMs, such as lasofoxefine [10, 11], and bazedoxefine [12]. However, one-third of the patients with ER+/PR+ tumors fail to respond to initial tamoxifen treatment, with many relapsing later [13, 14]. The development of tamoxifen resistance, studied predominantly in IDC is a complex phenomenon, involving an interplay of diverse cellular processes and signaling pathways, including upregulation of receptor tyrosine kinase activity leading to activation of ERK and PI3K pathway, altered expression of ERα and ERβ expression [15] and increased expression of miR-221/222 targeting p27/kip1 [16]. Recent studies demonstrate alteration of cellular metabolism as an important mechanism underlying development of drug resistance [17, 18, 19, 20]. Metabolic plasticity in cancer cells allows hijacking and remodeling existing metabolic pathways to foster cancer cell growth and survival impacting drug response [21]. For example, a recent study using tumor and adjacent normal tissues from IDC patients revealed distinct enrichment of one carbon metabolites in IDC tumors [22], yet similar metabolomic analysis of ILC cell lines and their endocrine resistant derivatives are limited.

To address these key gaps in ILC biology and therapeutic resistance, we developed and characterized tamoxifen-resistant ILC cell lines through concurrent transcriptional and metabolomic profiles with a goal to identify and characterize candidate therapeutic targets and biomarkers toward improved outcomes for ILC.

## Results

### Tamoxifen-resistant ILC cell lines

Tamoxifen-resistant (TAMR) cells were generated from two commercially available invasive lobular carcinoma (ILC) cell lines, MDA-MB-134-VI (MB134) and SUM44PE (SUM44). The MB134TAMR cells demonstrated marked morphological changes, with parental MB134 cells growing as a loosely adherent monolayer, while the MB134TAMR cells grown routinely in 100 nM 4-hydroxy tamoxifen (4-OHT) acquired a more cuboidal shape and formed larger adherent patches (**Fig. 1A**). SUM44 and SUM44TAMR cells had similar appearance growing as individually dispersed cells, with SUM44TAMR reflecting an increased proportion of cells with spindle like morphology, grown routinely in 500nM 4-OHT (increased concentration of 4-OHT due to inherent SUM44 tolerance).

**Figure 1.**
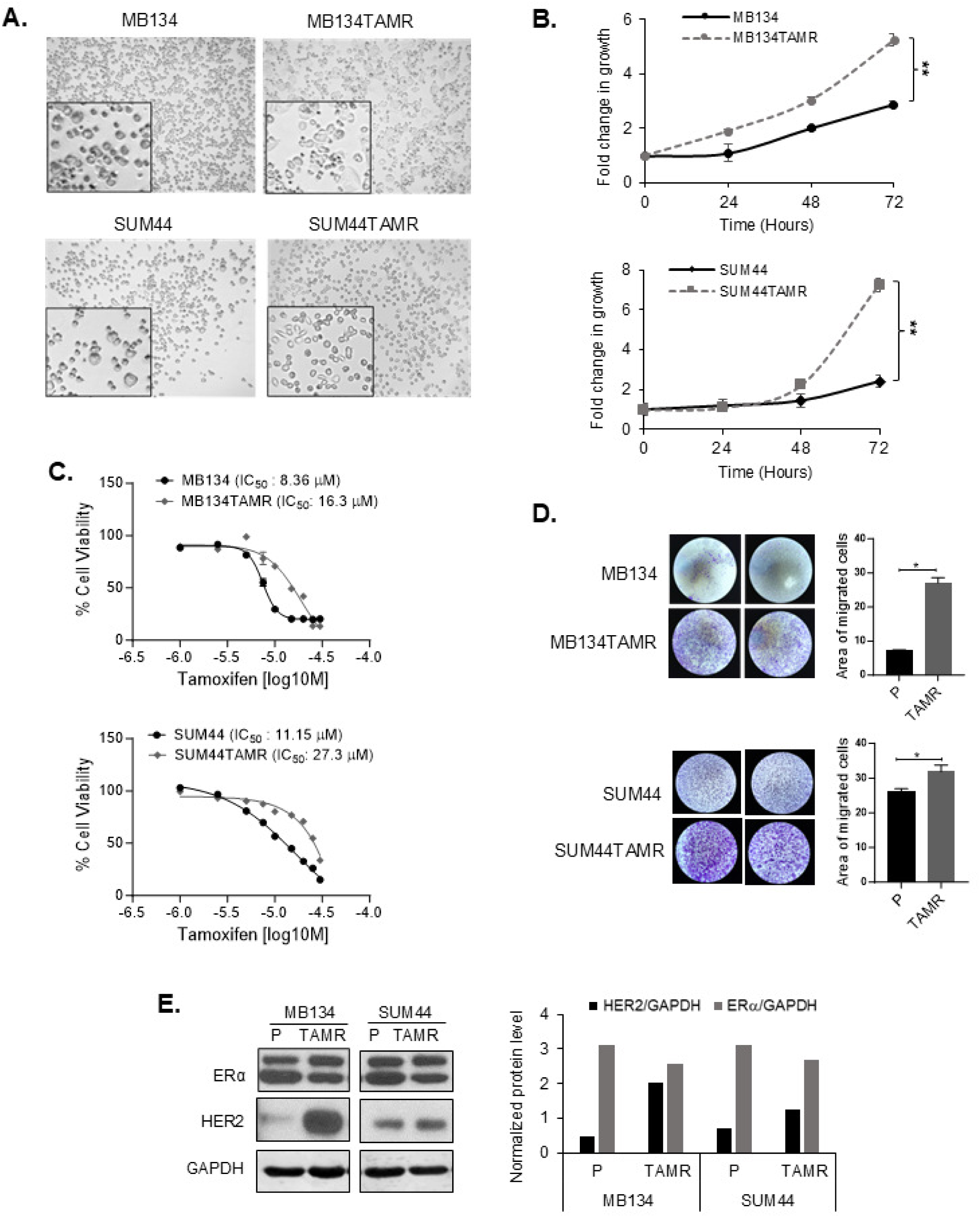
Characterization of Tamoxifen-Resistant ILC Cells. **A.** Phase contrast light microscopy images of parental and tamoxifen-resistant ILC cells with magnified images in the inserts. **B.** Growth kinetics of parental and tamoxifen-resistant MB134 (upper panel) and SUM44 (lower panel) cell lines. Fold change in growth normalized to day 0 at each time point over 72 hours. **C.** Dose response to tamoxifen in MB134 and MB134TAMR cells (upper panel) and SUM44 and SUM44TAMR cells (lower panel). Overnight cultures of exponentially growing cells were treated with vehicle or drugs for 5 days. IC_50_ for TAM was calculated using GraphPad Prism 10. **D.** Transwell Boyden chamber assays comparing the migratory capabilities of parental vs. TAMR, MB134 (upper panel) and SUM44 (lower panel) cell lines. Area covered by migrated cells were quantified in the bar diagram. **E.** Representative image of western blot and densitometry analysis of ERα and HER2 levels in parental vs. TAMR ILC cells. GAPDH was used as loading control. Representative of three independent experiments is presented in the figures. Statistical differences between groups were evaluated using Student’s t-test. Significance levels are indicated as follows: **p <0.001, *p <0.05. P= Parental, TAMR = Tamoxifen Resistant

Comparison of growth kinetics demonstrated significant increase in growth rate of TAMR cells relative to parental cells (MB134: *p = 0.008*, SUM44: *p = 0.005*) (**Fig. 1B**). In addition, TAMR cells showed >2-fold increase in the half-maximal inhibitory concentration (IC_50_) for tamoxifen compared to their respective parental counterparts (**Fig. 1C**), confirming functional tamoxifen resistance. Specifically, tamoxifen IC_50_ values were 8.4 μM and 16.3 μM for MB134 and MB134TAMR, respectively, and 11.15 μM and 27.3 μM for SUM44 and SUM44TAMR cells, respectively. Both TAMR cells demonstrated increased migration compared to the corresponding parental cells, with MB134TAMR 4-fold increase (*p=0.04,* **Fig. 1D**) and SUM44TAMR 1.2-fold increase relative to the corresponding parental cells (*p=0.046*, **Fig. 1D**). To evaluate if these changes were mediated through canonical breast cancer receptor expression, Western blot analysis revealed comparable ERα protein levels but TAMR cell lines expressed higher levels of HER2 than the parental lines (**Fig. 1E**): MB134-TAMR line showed a 4-fold increase and SUM44-TAMR had a 2-fold increase in HER2 expression.

Collectively, these data show that TAMR cells reflect morphological, phenotypic, and molecular changes relative to parental cell lines.

### Metabolic alterations associated with tamoxifen resistance in ILC cell lines

To elucidate the molecular changes associated with tamoxifen resistance in ILC, we subjected parental and TAMR derivatives of MB134 and SUM44 cell lines to metabolic profiling. Untargeted metabolomics profiling, involving comprehensive compound identification through databases like the Human Metabolite Database (HMDB) and an in-house high-resolution mass spectra database, was conducted on metabolites extracted from our representative cell lines. Using the polar metabolites fraction, we identified 120 metabolites across all samples with a QC coefficient of variation < 20%. Partial least square discriminant analysis (PLS-DA) was performed to summarize the overall metabolic differences between the cell phenotypes. Distinct and tight clustering of the experimental groups suggests distinct metabolic profiles pertaining to each cell line (**Supplementary Fig. S1A**). A heatmap was used to summarize the relative abundance of each metabolite across our queried cell lines (**Supplementary Fig. S1B**), where it became evident that the unique histological origin contributed to the metabolic uniqueness of each cell pair. Taking the abundance of unique metabolites in each pair of cell lines (parental *vs*. TAMR) into consideration, we chose to first analyze each pair separately. PLS-DA was thus performed to observe the metabolic differences of SUM44 parental vs. SUM44TAMR cells. **Fig. 2A** shows the independent clustering of SUM44 parental vs. SUM44TAMR cells, where model exhibited an R^2^ value of 0.99 and a Q^2^ score of 0.97, suggesting good model fit and predictive capability, respectively [23]. Our finding suggests that the acquired tamoxifen resistance contributes to a deregulated metabolic profile distinct from the parental line. Herein, a VIP plot highlights the top 15 deregulated metabolites driving separation of the PLS-DA model (**Fig. 2B**). Phosphocreatine, 5-amino levulinate and N-acetyl aspartic acid were the top deregulated metabolite across the SUM44 cell pairs. Importantly, many nucleotides and their derivatives, namely cytidine, uridine monophosphate, uridine diphosphate (UDP)-N-acetyl-D-galactosamine, and UDP-xylose were among the top deregulated metabolites driving separation of the PLS-DA model. The metabolic differences of the MB134 cell pair were also probed using a similar analysis, where a PLS-DA model demonstrated independent clustering of MB134 parental and MB134TAMR counterpart with an R^2^ value of 0.99 and a Q^2^ score of 0.95, indicating good model fit and predictive ability, respectively (**Fig. 2C**) [23]. The metabolites driving separation of the MB134 cell pair are depicted in the VIP plot in **Fig. 2D** where UDP-xylose, cytidine 5’-triphosphate and D-glucosamine 6-phosphate were identified as the top metabolites driving separation of the PLS-DA model. Collectively, our findings corroborate the notion that acquired resistance contributes to an aberrant metabolic profile.

**Figure 2.**
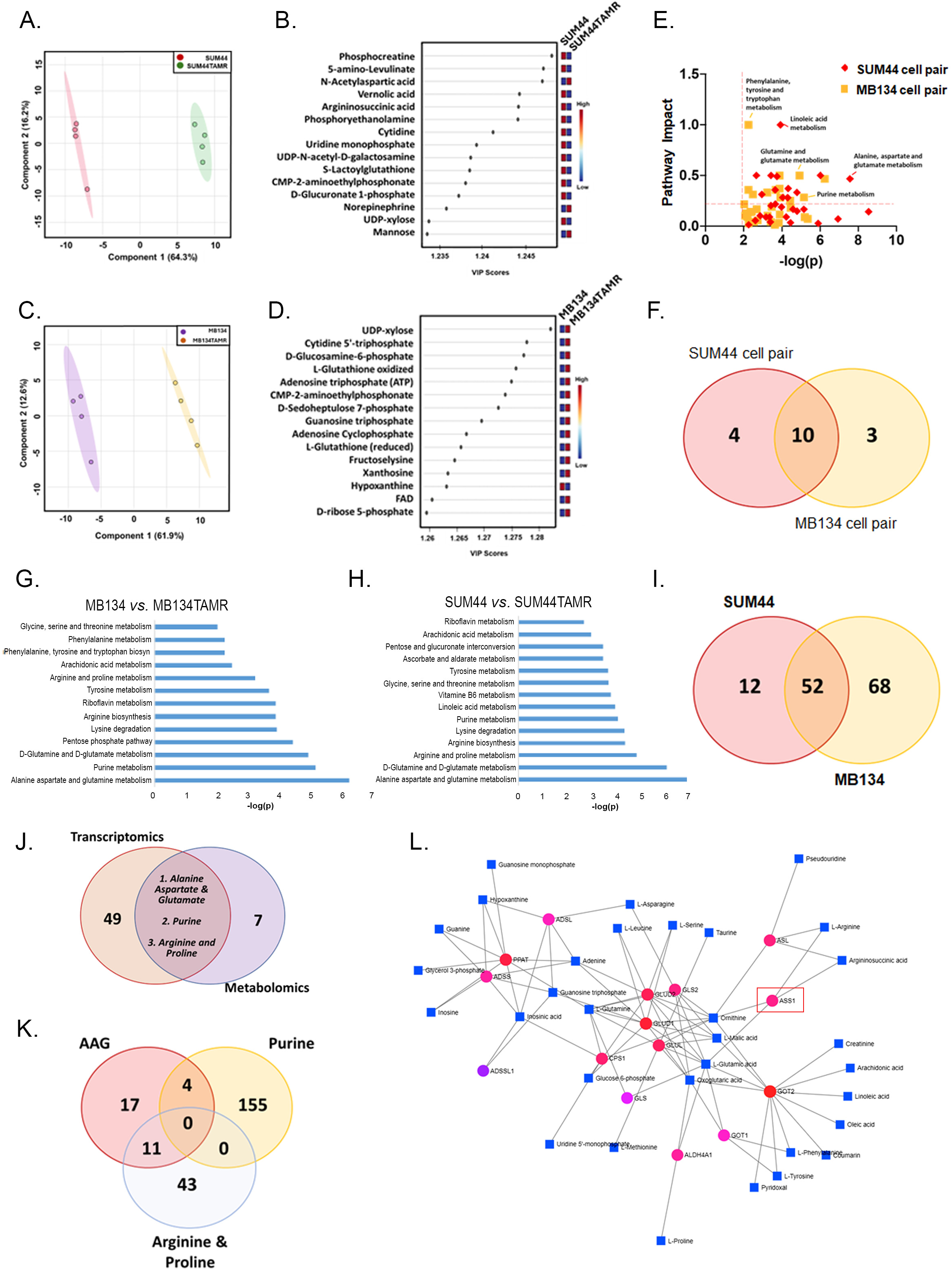
Metabolomic and transcriptomic analysis of paired ILC cell lines. **A.** Partial Least Squares Discriminant Analysis (PLS-DA) showing metabolic differences between parental and tamoxifen-resistant SUM44 cell pair. **B.** Variable Importance in Projection (VIP) plot highlighting the top 15 metabolites driving the separation between parental and resistant SUM44 cell pair. **C.** PLS-DA showing metabolic differences between parental and tamoxifen-resistant MB134 cell pair. **D.** VIP plot highlighting the top 15 metabolites driving the separation between parental and resistant MB134 cell pair. **E.** Overlap of altered pathways in parental vs. tamoxifen-resistant SUM44 and MB134 pairs, represented by impact scores. **F.** Venn diagram illustrating the number of shared and unique pathways altered in SUM44 and MB134 cell pairs, indicating common metabolic changes associated with tamoxifen resistance. **G.** Significantly deregulated KEGG pathways in parental vs. tamoxifen-resistant MB134 cells and, **H.** SUM44 cells as determined by RNA sequencing analysis of cell pairs in quadruplicate. **I.** Venn diagram showing the overlap of 52 deregulated pathways between parental and tamoxifen-resistant pairs of SUM44 and MB-134 cell lines. **J.** Venn diagram showing the overlap of metabolomic and transcriptomic data, identifying three mutually deregulated pathways in SUM44 and MB134 TAMR cells. **K.** Venn diagram showing the overlap of 15 genes within the three mutually deregulated pathways. **L.** Gene-metabolite interaction map illustrating interactions between three deregulated pathways. Circles represent genes and squares represent metabolites.

Systemically, quantitative enrichment analysis uncovered 14 significantly altered pathways that were associated with the acquired tamoxifen resistance in our SUM44 cell pair [–log(p) value > and pathway impact > 0.2]. In comparison, the analysis uncovered 13 significantly altered pathways that were associated with the acquired tamoxifen resistance in the MB134 cell pair. The pathways included alanine, aspartate and glutamate metabolism, tyrosine metabolism, glycine, linoleic acid metabolism, purine metabolism, lysine degradation, arginine biosynthesis, arginine and proline metabolism, as well as D-glutamine and D-glutamate metabolism (**Fig. 2E**). A full list of the significantly deregulated pathways between our queried cell pairs is summarized in **Supplementary Table 1.** To investigate mutually deregulated metabolic processes driven by acquired tamoxifen resistance, we identified 10 mutually deregulated pathways between the TAMR and parental cell lines (**Fig. 2F & Table 1**). The pathways deregulated in both the cell lines were predominantly amino acid metabolism pathways such as alanine, aspartate and glutamate metabolism, arginine biosynthesis and lysine degradation, but also included nucleotide metabolic processes such as purine metabolism pathway. Importantly, identification of <u>shared</u> metabolic changes across both ILC cell pairs identifies the prominent metabolic processes consistently altered as a result of acquired tamoxifen resistance.

**Table 1:** List of 10 mutually deregulated pathways between the parental and TAMR cell lines revealed by metabolomic analysis.

| **Pathway Name** | **-log(p)** | **Impact** | **Cell pair** |
| --- | --- | --- | --- |
| Alanine, aspartate and glutamate metabolism | 7.57<br>6.24 | 0.47<br>0.47 | SUM44<br>MB134 |
| Arginine biosynthesis | 4.33<br>3.88 | 0.37<br>0.37 | SUM44<br>MB134 |
| Lysine degradation | 4.30<br>3.92 | 0.28<br>0.28 | SUM44<br>MB134 |
| Arachidonic acid metabolism | 2.95<br>2.48 | 0.31<br>0.31 | SUM44<br>MB134 |
| D-Glutamine and D-glutamate metabolism | 6.02<br>4.93 | 0.50<br>0.50 | SUM44<br>MB134 |
| Riboflavin metabolism | 2.66<br>3.88 | 0.50<br>0.50 | SUM44<br>MB134 |
| Glycine, serine and threonine metabolism | 3.66<br>2.02 | 0.22<br>0.22 | SUM44<br>MB134 |
| Purine metabolism | 4.04<br>5.16 | 0.28<br>0.28 | SUM44<br>MB134 |
| Arginine and proline metabolism | 4.78<br>3.23 | 0.33<br>0.33 | SUM44<br>MB134 |
| Tyrosine metabolism | 3.64<br>3.68 | 0.36<br>0.36 | SUM44<br>MB134 |

### Altered nucleic acid and amino acid pathways associated with tamoxifen resistance

To further validate the observed metabolic alterations and delineate the underlying metabolic rewiring, we subjected all four cell lines to RNAseq transcriptomic analysis. The volcano plots show the differentially expressed genes between the SUM44 pair (**Supplementary Fig.S2A**) and MB134 pair (parental vs. TAMR) (**Supplementary Fig.S2B**). The genes deregulated in our RNA-seq data were subjected to Gene Set Enrichment analysis (GSEA). A total of 30,000 genes were analyzed and gene sets were mapped to KEGG database. In line with our metabolomics analysis, transcriptomic data was analyzed separately for each pair of cell lines. Significantly altered pathways were identified with a cut-off of FDR < 0.2 for significance for both MB134 cell pairs and SUM44 cell pairs where up to top 14 pathways are shown in **Fig. 2G** and **Fig. 2H**, respectively. Mutually altered pathways across both cell pairs were next analyzed, with 52 pathways identified to be significantly deregulated as a result of acquired tamoxifen resistance (**Fig. 2I**). A subset of the pathways deregulated in both MB134TAMR and SUM44TAMR cells is highlighted in **Table 2**.

**Table 2:**
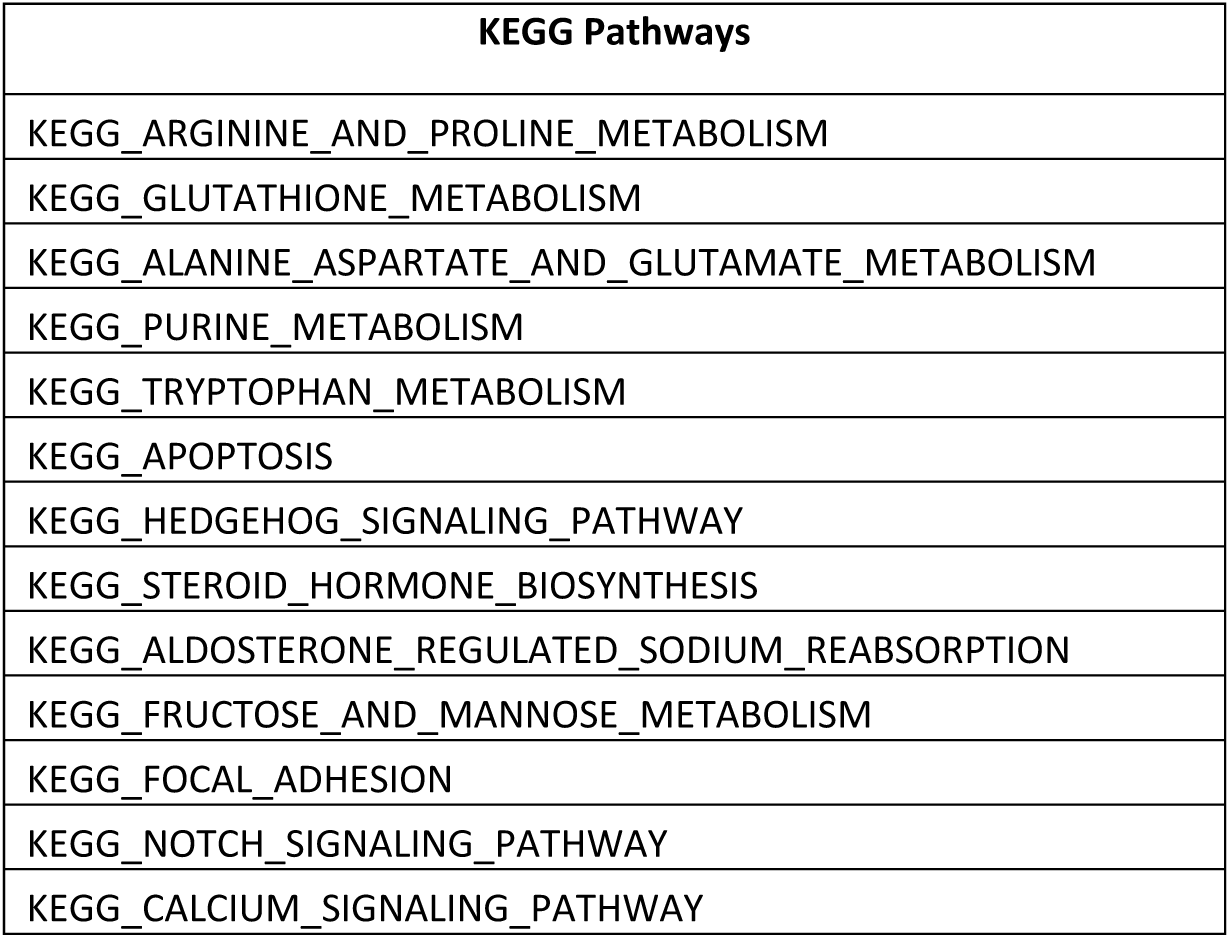
Subset of deregulated pathways significantly altered in both ILC cell pairs as revealed by transcriptomic analysis.

To further enable mechanistic understanding, an integrated analysis of both metabolomics and transcriptomics revealed that only three mutually deregulated metabolic pathways exhibited similar alterations in both parental and TAMR ILC cells (across both of our metabolomics and transcriptomics datasets): 1) alanine, aspartate, and glutamate metabolism (AAG); 2) purine metabolism; and 3) arginine and proline metabolism (**Fig. 2J**). To further investigate these metabolic changes and to confirm the observed trend in tamoxifen resistant cells, we identified genes overlapping in all dysregulated pathways. While a single gene was not at the intersection of all three pathways, 15 genes were found to partake in at least 2 unique metabolic processes **Fig. 2K**. There were 4 gene overlap between AAG and Purine metabolism pathway, namely *ADSS1*, *ADSS2*, *ADSL* and *PPAT*. Additionally, there were 11 genes that overlapped between AAG and Arginine & proline metabolism pathways, namely, *ALDH4A1*, *ASS1*, *ASL*, *CPS1*, *GLS*, *GLS2*, *GLUL*, *GLUD1*, *GLULD2*, *GOT1*, *GOT2* (**Supplementary Table 2**). As these genes partake in an intricate metabolic network, a gene metabolite interaction map between our three deregulated pathways was mapped in **Fig. 2L** to investigate potential association with tamoxifen resistance. This analysis provides a comprehensive view of the molecular landscape associated with tamoxifen resistance in ILC cell lines, with *ASS1 (Arginosuccinate Synthase 1)* emerging as one of the central players in the observed metabolic and transcriptomic alterations connecting amino acid and nucleotide biosynthesis. Collectively, the integration of our unbiased analyses was able to identify 3 unique metabolic pathways directly associated with tamoxifen resistance at both the metabolite and transcript level. Overlap of the genes within the deregulated pathways identified 15 mutually expressed genes across at least 2 pathways, where *ASS1* was determined to be one of the central metabolic hubs within this aberrant metabolic network. ASS1 is a key enzyme in urea cycle, synthesizing arginosuccinate from aspartate and citrulline. Downregulation of ASS1 could lead to metabolic diversion of aspartate towards nucleic acid biosynthesis fostering cell proliferation.

### Methylation-mediated downregulation of *ASS1* associated with tamoxifen resistance in ILC cell lines

Transcriptomics data revealed a significant down-regulation of *ASS1* expression in both MB134-TAMR (95%, *p= 0.01)* and SUM44-TAMR cells (90%, *p=0.019)*; (**Fig. 3A**). This finding was further confirmed in three independent biological replicates by qRT-PCR, where consistent decrease in *ASS1* mRNA levels in TAMR cells (MB134-TAMR: 40%, *p<0.0001* and SUM44-TAMR: 73%, *p<0.0001*) compared to the respective parental cells were observed (**Fig. 3B**). We also evaluated *ASS1* expression in ILC cells grown long-term in estrogen deprived (LTED) medium. The MB134-LTED and SUM44-LTED cells grown in absence of estrogen demonstrated resistance to tamoxifen [24]. Interestingly, two different clones of LTED cells derived from each ILC cell lines showed a significant decrease in *ASS1* expression (MB134-LTED: >90%, *p<0.0001*, SUM44-LTED: 90%, *p<0.0001*; **Fig. 3B**). Marked reduction of ASS1 protein (60-90%) was observed in TAMR and LTED cells compared to the corresponding parental cell lines (all *p<0.05*, **Fig. 3C**). This observation raises the possibility that estrogen may play a regulatory role in modulating *ASS1* expression, although comparable ERα protein levels (as shown in **Fig 1C**).

**Figure 3.**
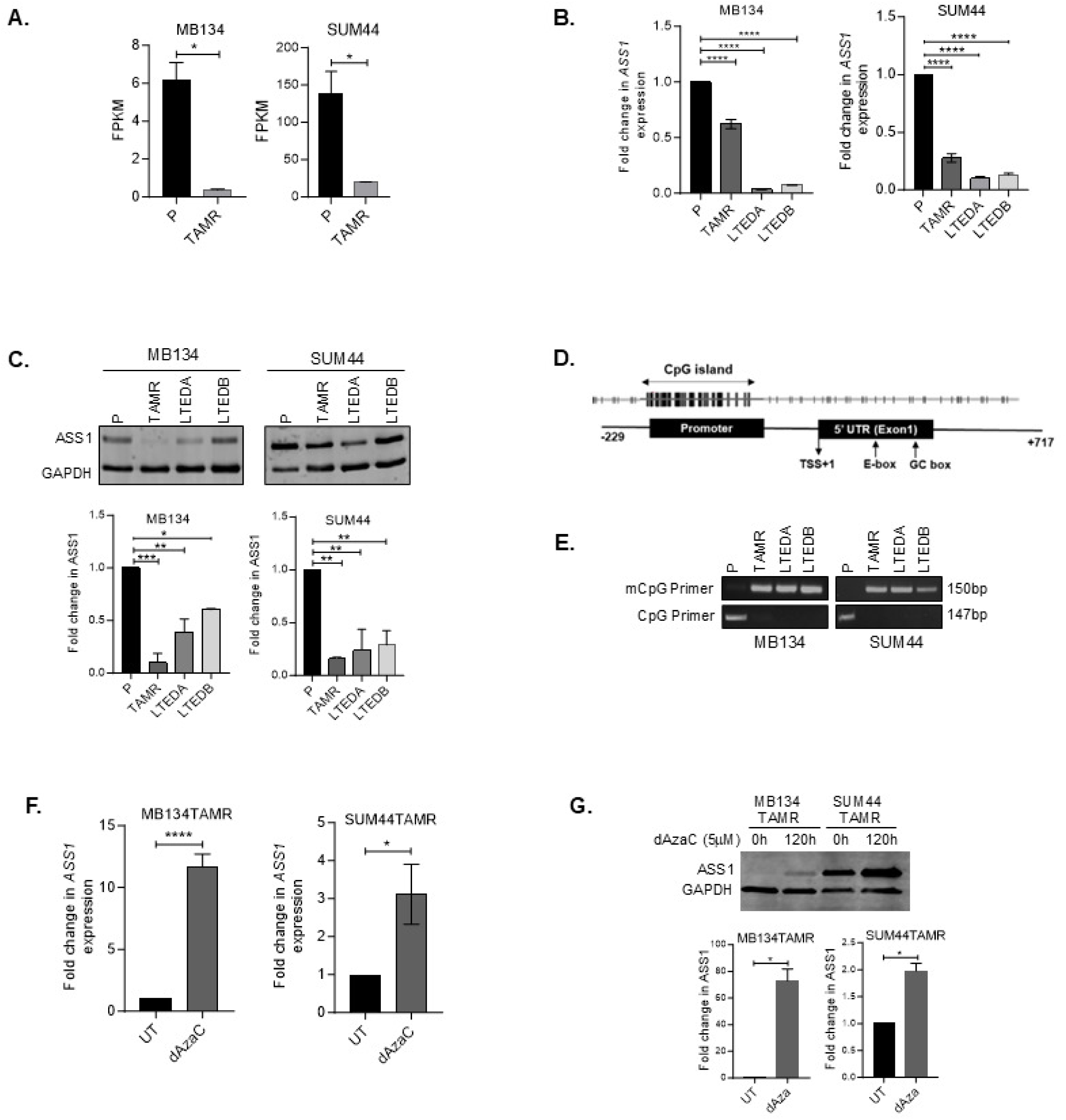
Downregulation of ASS1 in ILC-TAMR and LTED Cells. **A.** Comparative expression of *ASS1* in parental vs. parental and tamoxifen-resistant ILC cell lines as determined by change in Fragments Per Kilobase of transcript per Million mapped reads (FPKM) obtained from RNA seq data. **B.** qRT-PCR analysis showing *ASS1* expression, and **C.** western blot and densitometry analysis showing ASS1 protein levels in MB134 and SUM44 cells, their respective tamoxifen resistant and LTED derivatives (LTEDA and LTEDB). **D.** Schematic diagram showing the CpG island in the *ASS1* promoter. **E.** Analysis of the *ASS1* promoter region using MS-PCR showing amplification of 150 bp methylated DNA in TAMR and LTED derivatives of ILC cell lines and 147 bp unmethylated DNA in parental ILC cell lines. **F.** qRT-PCR analysis showing *ASS1* expression in MB134TAMR and SUM44TAMR cells that are either left untreated (UT) or treated with 5 μM 5-Aza-2’-deoxycytidine (dAzaC) for 120 hours. **G.** Western blot and densitometry analysis showing ASS1 protein levels in MB134TAMR and SUM44TAMR cells after 120 hours of dAzaC treatment (n=2 experiments). Error bars represent the standard deviation of triplicates. Significance levels are indicated as follows: ****p <0.0001, ***p <0.001, **p <0.01, *p <0.05.

We next investigated the mechanism underlying silencing of *ASS1* in TAMR ILC cells. Computational analysis revealed presence of a CpG island in the 5′ regulatory region of the *ASS1* gene spanning from −499 bp to −6bp with +1 as the transcription start site (**Fig 3D**). Methylation-specific PCR (MSP) using methyl CpG specific primers led to amplification of a 150bp product from TAMR and LTED cell DNA but not from parental cell DNA. Use of primers specific for unmethylated DNA led to amplification of a147 bp product from parental cell DNA, but not from TAMR and LTED cell DNA, (**Fig. 3E**). Our data demonstrates methylation of the *ASS1* promoter in TAMR and LTED cells but not in the parental counterparts.

To determine if treatment with a demethylating agent would reverse methylation at this specific 5′ regulatory region of the *ASS1* gene and enhance expression of *ASS1,* TAMR cells were treated with established demethylating agent decitabine (5-Aza-2’-Deoxycytidine, dAzaC) at 5 μM. When treated for 120 hours, MB134-TAMR cells showed a 12-fold increase (*p< 0.0001*), and SUM44-TAMR cells showed a 3-fold increase (*p=0.01*) in *ASS1* mRNA, when compared to untreated cells (**Fig. 3F**). Similarly, a 72-fold and ∼2-fold increase of ASS1 protein in dAzaC treated MB134-TAMR (*p=0.0078*) and SUM44-TAMR (*p=0.01*) cells respectively were observed (**Fig. 3G**). These data support the notion that methylation of the CpG island in *ASS1* promoter contributes to its downregulation in tamoxifen resistant cells.

To determine if *ASS1* downregulation is sufficient to mediate tamoxifen resistance, we knocked down *ASS1* in MB134 and SUM44 cells using shRNA (*shASS1* cells). In MB134-*shASS1* cells there was a 50% reduction in ASS1 mRNA (*p=0.0012),* and protein was barely detectable (*p=0.0003)*, whereas in SUM44-*shASS1* cells 80% reduction in both mRNA (*p<0.0001)* and protein (*p=0.0001)* levels were observed (**Fig. 4A** & **4B**). The *shASS1* cells demonstrated increased tolerance to tamoxifen as evidenced by increase in IC_50_ values for tamoxifen compared to the parental cells. For MB134 cells, IC_50_ values are 9.0 µM *vs.* 15.2 µM in pLKO vs. *shASS1* cells (p*=0.017*), and for SUM44 cells the values are 11.9 µM *vs.* 18.4 µM in pLKO vs. *shASS1* cells (*p=0.05*) (**Fig. 4C**). In addition, we have observed a small but significant increase in growth rate when *ASS1* is knocked down in the ILC cell lines (**Fig. 4D**, MB134: pLKO vs. *shASS1, p=0.04* and SUM44: pLKO vs. *shASS1, p=0.03*). These findings suggest that reduced expression of *ASS1* promotes cell growth, and resistance to tamoxifen.

**Figure 4.**
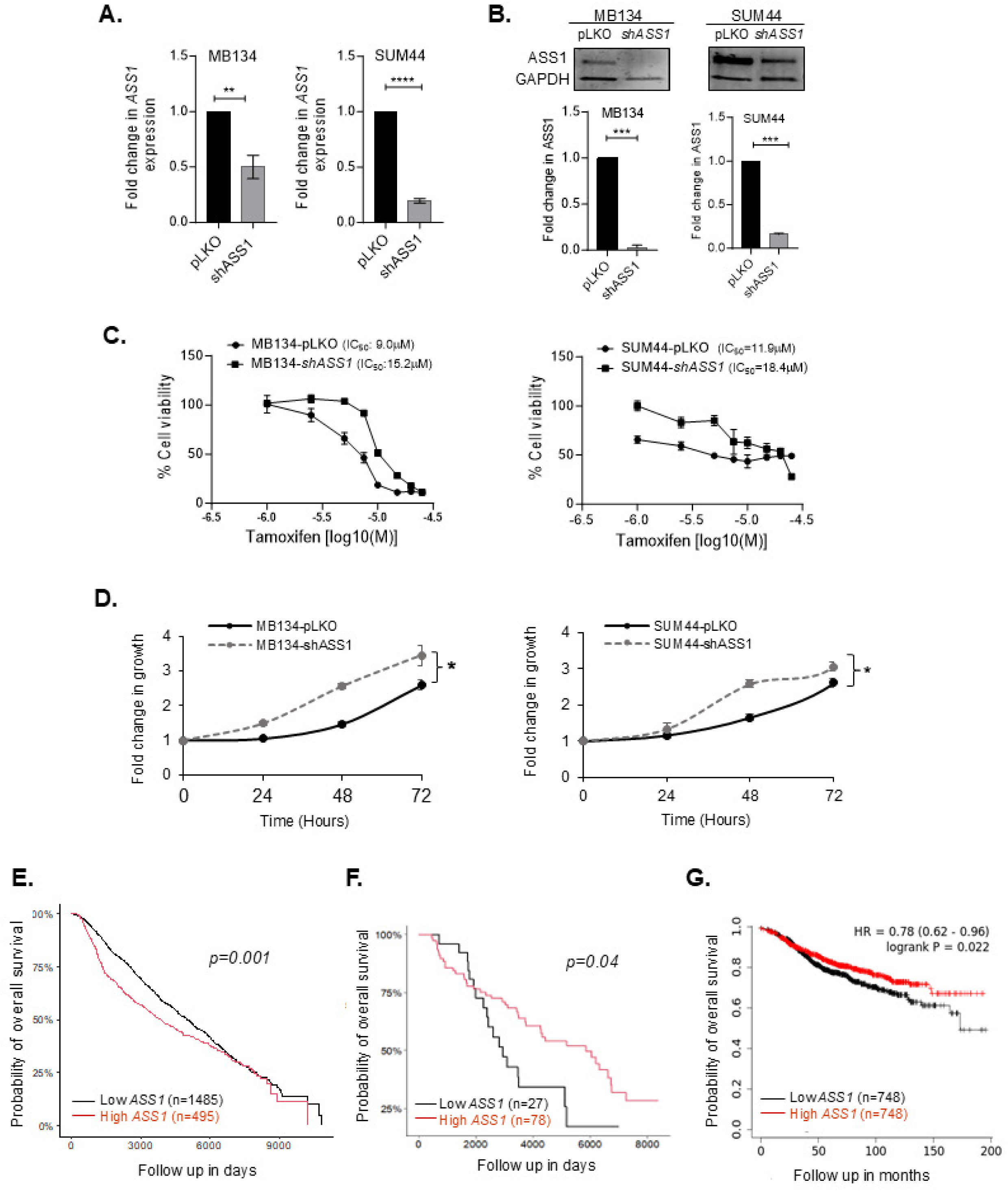
*ASS1* knockdown in ILC cells. MB134 and SUM44 cells were transduced with plasmids expressing shRNA targeting *ASS1* or the corresponding empty vector to produce *shASS1* and control (pLKO) cell lines. **A.** qRT-PCR analysis showing *ASS1* expression, and **B.** western blot and densitometry analysis of ASS1 protein levels in *ASS1* knockdown (*shASS1*) vs. pLKO (control) MB134 and SUM44 cell lines. **C.** Dose response to tamoxifen and IC_50_ of tamoxifen in *ASS1* knockdown (*shASS1*) vs. pLKO (control) MB134 (left panel) and SUM44 cell lines (right panel). **D**. Comparison of growth kinetics of *shASS1* vs. pLKO derivatives of MB134 (left panel) and SUM44 cell lines (right panel). Fold change in growth is normalized to day 0 at each time point over 72 hours. **E.** Overall survival analysis of all breast cancer patients in relation to *ASS1* expression using METABRIC data set. **F**. Overall survival analysis of ILC patients who received endocrine therapy in relation to *ASS1* expression using METABRIC data set. **G.** Overall survival analysis of breast cancer patients who received endocrine therapy in relation to *ASS1* expression using K-M plotter dataset. ****p <0.0001, ***p <0.001, **p <0.01, *p <0.05.

To determine the significance of *ASS1* downregulation in breast cancer patients with respect to disease outcome, we used publicly available datasets. Using the METABRIC dataset, when all subtypes are included, we found that upregulation – not downregulation - of *ASS1* (upper quartile) is prognostic of poor overall survival (OS) in the breast cancer population (**Fig.4E**, *p=0.001*), suggesting our findings are not generalizable to all breast cancer subtypes together. When we specifically analyzed ILC patients treated with endocrine therapy (the focus of this study), low *ASS1* expression (lower quartile) is prognostic of poor OS, and the correlation was significant with a *p-value* of 0.04 (**Fig. 4F**). We did evaluate all endocrine-treated breast cancer patients in METABRIC, and *ASS1* was not predictive of OS (data not shown). Thus, there is evidence that low *ASS1* expression, associated with tamoxifen resistance experimentally, is associated with worse survival among endocrine therapy-treated patients and, specifically, endocrine therapy-treated patients with ILC. To evaluate an orthogonal dataset, we used the K-M Plotter dataset to analyze the correlation between *ASS1* expression and OS of breast cancer patients who received endocrine therapy and observed that lower ASS1 expression was associated with worse OS (**Fig 4G**, *p=0.022*).

### Nucleic acid biosynthesis and pathway intermediates in tamoxifen resistant ILC cells

Aberrant metabolism of amino acids and one-carbon units in cancers are acknowledged contributors to nucleic acid synthesis, fostering proliferative signaling, resistance to cell death, and metastasis [25]. Purine metabolism is one of the three pathways that was found to be mutually deregulated in both TAMR cells in our multi-omics analysis (**Fig.2J**). When subjected to metabolomic analysis, *ASS1* knock down and the respective control cells (*shASS1* vs. pLKO) derived from MB134 and SUM44 cell lines showed independent clustering (**Supplementary Fig. S3A and S3C**). The VIP plots highlight the top metabolites driving the separation of the *ASS1* knock down *vs.* the control cells (**Supplementary Fig. S3B and S3D)**. Importantly, similar enrichment of purine metabolism was observed in both MB134 (*p=0.003)* and SUM44 (*p=0.0008)* cells upon *ASS1* knock down (**Fig. 5A & 5B**). In addition, the observed enrichment of alanine, aspartate and glutamate metabolism in SUM44-*shASS1* cells, suggests association between *ASS1* downregulation and metabolic pathways altered in TAMR cells. Our untargeted metabolomics analysis uncovered changes in intermediates of nucleotide metabolism upon acquiring tamoxifen resistance. In particular, we noticed an increased in intracellular abundance of the purine intermediates deoxyguanosine-monophosphate (dGMP) (*p=0.03*), guanosine diphosphate (GDP) (*p=0.03*) and guanosine monophosphate (GMP) (*p=0.02*) (**Fig.5C**). Additionally, an increase in the pyrimidine intermediate cytosine monophosphate (CMP) was also evident in the MB134TAMR cells (*p=0.002*) (**Fig.5D**). Similarly, in SUM44TAMR cells, we observed small but significant increase in ADP (*p=0.009*) and ATP (*p=0.05*), as well as an elevation in cytosine level (*p= 0.0006*) (**Fig. 5E & 5F**).

**Figure 5.**
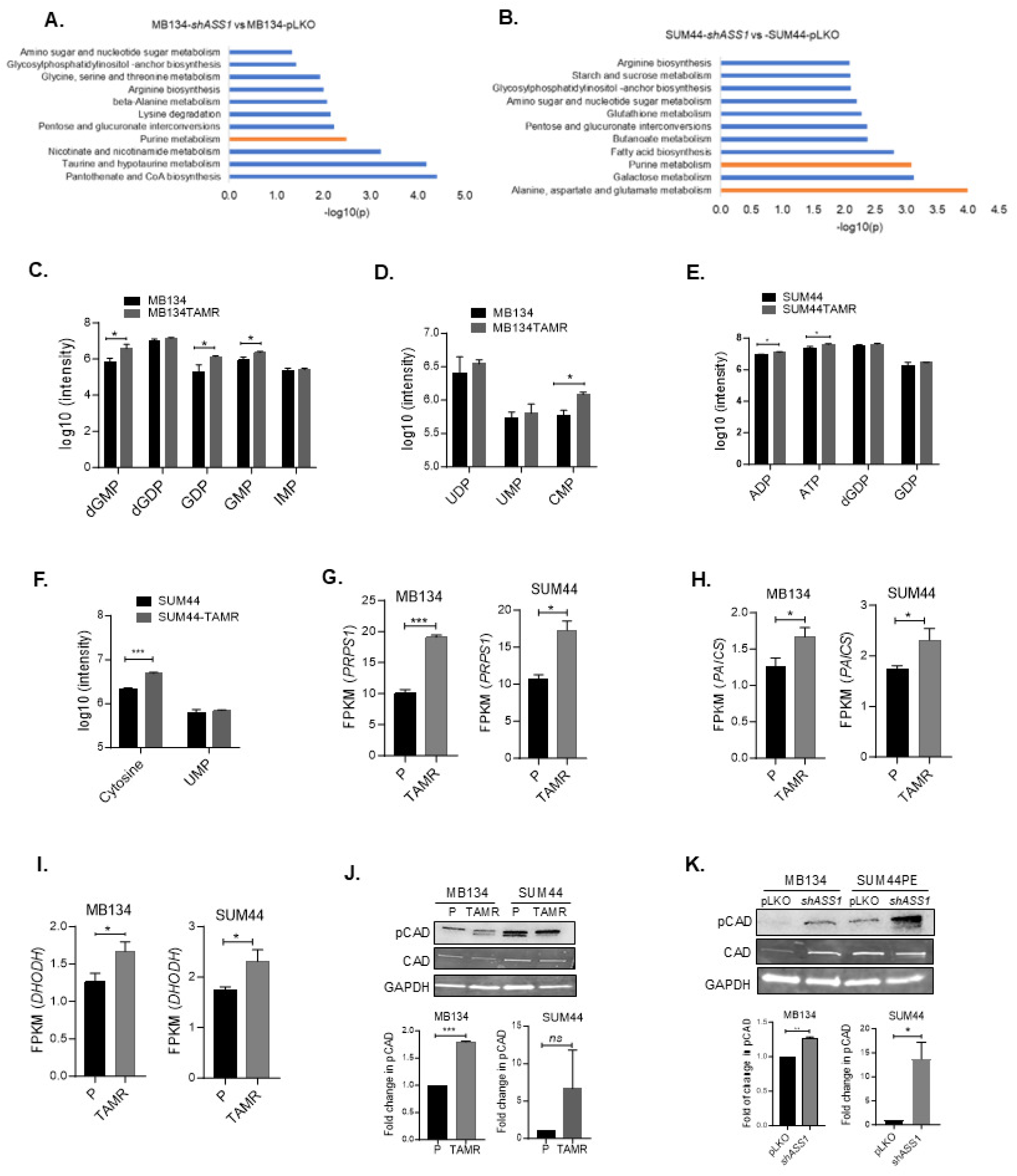
Nucleic acid biosynthesis and pathway intermediates in tamoxifen resistant ILC cells. **A.** Significantly enriched metabolic pathways in *shASS1* vs. pLKO derivatives of MB134, and **B.** SUM44 cell lines. **C.** Changes in the levels of purine (dGMP, dGDP, GDP, GMP, IMP) and pyrimidine (UDP, UMP and CMP) nucleotides in parental (P) vs. tamoxifen-resistant (TAMR) MB134 cells. **E.** Changes in the levels of purine (ATP and ADP, dGTP, GDP) and **F.** pyrimidine (Cytosine, UMP) nucleotides in parental (P) vs. tamoxifen-resistant (TAMR) SUM44 cells. **G.** Expression levels of *PRPS1*, **H.** *PAICS,* and **I.** *DHODH* in MB134 and SUM44 cell pairs as analyzed from RNA seq data. (FPKM: Fragments Per Kilobase Per Million reads). **J.** Representative picture of pCAD^S1859^ (pCAD) levels in parental and TAMR cells and **K.** in control (pLKO) vs. *shASS1* derivatives of MB134 and SUM 44 cell pairs. Bar diagrams show average of more than one independent experiments (n=2). ***p <0.001, **p <0.01, *p <0.05.

Using the transcriptomics data we next investigated if expression of additional genes involved in purine and pyrimidine synthesis pathways are altered in TAMR cells. We observed significant upregulation of *PRPS1* (*phosphoribosyl pyrophosphate synthetase 1*) in MB134TAMR (*p=0.0008*) and SUM44TAMR cells (*p=0.017*) when compared to the respective parental cells (**Fig. 5G**). Phosphoribosyl pyrophosphate is essential for both *de novo* and salvage pathway of purine and pyrimidine biosynthesis. Additionally, we found *PAICS (phosphoribosylaminoimidazole carboxylase and phosphoribosylaminoimidazolesuccincarboxamide synthase)* converting carboyaminoimidazole ribonucleotide (CAIR) to N-Succinocarboxyamide-5-aminoimidazole ribonucleotide (SAICAR), an intermediate of purine biosynthesis to be significantly upregulated in both MB134TAMR (*p=0.015*) and SUM44TAMR cells (*p=0.01*) (**Fig. 5H**). Furthermore, there is significant increase in *DHODH* (*dihydroorotase dehydrogenase*) converting dihydroorotate to orotate, a pyrimidine biosynthesis intermediate in both MB134TAMR (*p=0.015*) and SUM44TAMR cells (*p=0.01*) (**Fig. 5I**). *ADSS2 (adenylosuccinate synthase 2),* a crucial enzyme for converting IMP to AMP is upregulated significantly in MB134TAMR cells (*p=0.036*) but not in SUM44TAMR cells (**Supplementary Fig. S4A and S4B**). Collectively, multiple enzymes in the nucleotide biosynthesis pathways are upregulated in the TAMR cells.

CAD (Carbamoyl-phosphate synthase 2, Aspartate transcarbamoylase, Dihydroorotase), a key trifunctional enzyme utilizes aspartate as substrate and executes the first three steps of pyrimidine biosynthesis. Phosphorylation at Ser1859 leads to activation of CAD. To this end we have evaluated pCAD^S1859^ level in parental and TAMR cells, where a 1.8-fold increase in pCAD was observed in MB134TAMR compared to MB134 cells (**Fig. 5J**, *p=0.0003*). SUM44TAMR cells showed a trend toward increase in pCAD, compared to SUM44 cells. When analyzed in *ASS1* knockdown cells, significant increase in pCAD^S1859^ levels was observed in both MB134-*shASS1 (1.3-fold, p=0.003)* and SUM44-*shASS1 (11-fold, p= 0.036)* cells compared to the control cells (**Fig. 5K**).

In conclusion, multiple cellular pathways are affected in the course of acquiring resistance to anti-estrogen that promotes proliferation of the resistant cells.

### Therapeutic targeting of *ASS1* methylation and nucleotide biosynthesis to overcome tamoxifen resistance

Our data demonstrated methylation mediated silencing of *ASS1* that correlates with poor outcome in endocrine treated ILC patients. We rationalized that increased expression of *ASS1* by promoter demethylation will improve efficacy of tamoxifen. To this end TAMR ILC cells were pre-treated with 5 µM dAzaC for 120 hours, untreated cells were used as control. Subsequently, MTT assays were performed with both dAzaC pre-treated and untreated cells, exposing them to increasing concentrations of tamoxifen. Our data showed significantly higher inhibition of dAzaC pre-treated MB134TAMR when treated with 7.5 and 10 µM tamoxifen when compared to the effects on dAzaC -untreated cells (*p=0.04* and *0.03,* respectively. **Fig. 6A**). Similarly, SUM44TAMR cells were significantly more sensitive to 10 µM tamoxifen when compared with untreated cells (*p=0.05*, **Fig. 6B**), suggesting this to be a potential therapeutic combination for ILC patients with acquired resistance to endocrine therapy.

**Figure 6.**
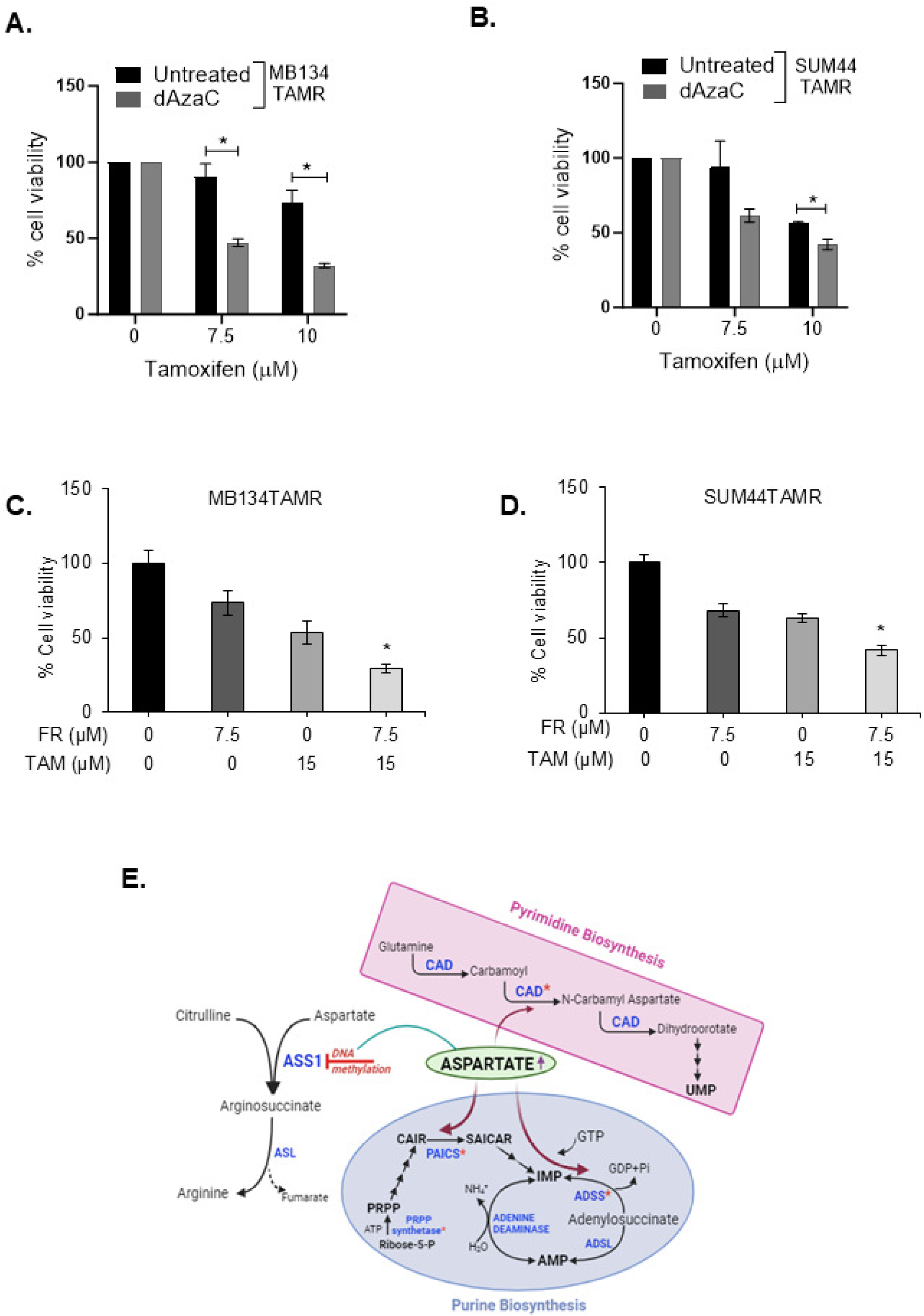
Combined treatment of tamoxifen-resistant ILC cell lines with tamoxifen and drugs targeting *ASS1* methylation or nucleotide biosynthesis. **A.** MB134TAMR and **B.** SUM44TAMR cells were pretreated with 5μM dAzaC for 120 hours. Untreated (vehicle) cells were used as control. Viability of dAzaC treated and control cells in presence of indicated doses of tamoxifen was compared after additional 5 days of treatment. **C.** Comparison of cell viability after treatment of MB134TAMR and **D.** SUM44TAMR cells with 10 μM Farudodstat (FR) alone and in combination with 15 μM tamoxifen for 5 days. Data is representative of three independent experiments conducted in triplicate for each treatment condition. **E.** Schematic diagram showing how methylation mediated silencing of *ASS1* could lead to augmentation of purine and pyrimidine biosynthesis pathway in tamoxifen-resistant ILC cells, where the enzymes that uses aspartate as substrate are highlighting. Statistical significance indicated as *p < 0.05.

Based on the observation that multiple pyrimidine biosynthesis intermediates are deregulated in TAMR ILC cells, we next examined if inhibition of *de novo* pyrimidine biosynthesis is an additional avenue to improve tamoxifen efficacy in endocrine resistant ILC. Farudodstat (ASLAN003) is an FDA-approved inhibitor of DHODH, a rate-limiting enzyme for *de novo* pyrimidine biosynthesis. Our data showed increased *DHODH* expression in TAMR cells (**Fig.5I**). To test the combinatorial effect of tamoxifen and farudodstat, TAMR cells were treated with farudodstat alone or in combination with tamoxifen for 120 hours. Combined treatment of MB134TAMR cells with 15 μM tamoxifen and 7.5μM farudodstat led to an 71% inhibition of cell viability. In comparison, treatment with 15 µM tamoxifen alone resulted in a 47% reduction, and 7.5 µM farudodstat alone led to a 26% reduction in cell viability. These results indicate a synergistic effect of the combined treatment, as reflected by a combination index (CI) of 0.85931 *(p= 0.02*, **Fig.6C**). In SUM44TAMR cells, the combination resulted in synergistic inhibition as well, where a 58% inhibition in cell viability was observed (CI=0.84613), compared to 37% and 32% inhibition by tamoxifen alone and farudodstat alone respectively (*p=0.05,* **Fig. 6D**). This data suggests that combined targeting of nucleic acid biosynthesis and estrogen signaling could be a potential therapeutic option for ILC patients with acquired resistance to tamoxifen.

## Discussion

One of the corner stones in treating patients with ER+ ILC is endocrine therapy [13, 26]. In general, ILC tumors have lower response rates to chemotherapy because of low proliferation index. Resistance to endocrine therapy poses a major challenge in managing the disease effectively. Up to 40% of ER+ breast cancer patients’ tumors may develop tamoxifen resistance during the initial phase of treatment, with an additional 25% developing resistance over time [27]. Although the mechanisms underlying endocrine resistance have been studied extensively in ER+ IDC (reviewed by Osborne *et.al.* [28]), these two subtypes of breast cancer are distinct in terms of histo-morphology, disease progression, recurrence, and outcome, underscoring the importance of studying endocrine resistance <u>specifically</u> in ILC. This is the first study to our knowledge where a multi-omics approach was used to compare tamoxifen resistant ILC cell lines with their parental counterpart and identify aberrations in amino acid and nucleotide biosynthesis pathways in the resistant cells. In these cells, acquired resistance to tamoxifen resulted in down regulation of *ASS1*, a key enzyme at the intersection of these pathways, and corelates with poor overall survival of ILC patients, specifically those who received endocrine therapy.

Our recent review highlights the studies that investigated mechanism of endocrine resistance in ILC [13]. Reduced expression of *ERα*, increased expression of estrogen-related receptor γ (*ERRγ*) [29], activation of AP1-dependent transcription [29, 30], frequent mutation of *PTEN* and *PIK3CA* [31], activation of SREBP1 driving lipid and cholesterol metabolism specifically in resistance to aromatase inhibitors [32, 33], and the involvement of WNT4 in estrogen-induced growth [34], has been associated with endocrine resistance in ILC. Additionally, mutations in *FOXA1*, a pioneer factor for ER-mediated transcription, confers endocrine resistance by increasing *FOXA1* expression and activity [35].

The major challenges in studying ILC is lack of established and authentic cell lines along with slow growth rate of the tumor reflected in the mouse models of ILC, particularly in orthotopic models [36]. A recent study has systematically analyzed several ILC/ILC like cell lines and identified additional cell lines harboring key molecular features of ILC [37]. These cell lines are promising for future studies. Currently, the two most commonly used and universally accepted cell lines to study ILC are MDA-MB-134-VI and SUM44PE, confirming the relevance to ILC biology and used in this study. Both these cell lines are ER+ and lack E-cadherin, harboring the most common features of ILC tumors. SUM44PE was isolated from a patient refractile to endocrine therapy and is therefore *de novo* resistant to endocrine therapy [38], whereas MB134 cells were isolated from pleural fluid of a patient diagnosed with papillary mammary carcinoma, later classified as luminal subtype [36, 39]. MB134TAMR cells generated in our lab models acquired anti-estrogen resistance. Increased tolerance of the SUM44TAMR cells to tamoxifen is not therefore expected to fully mimic the characteristics observed in MB134TAMR cells. This poses additional challenge and limits the ability to study and validate the mechanisms underlying development of anti-estrogen resistance in ILC patients. However, long term exposure of the two ILC cell lines to tamoxifen, the classic estrogen receptor modulator to generate TAMR cells and to estrogen deprivation to generate LTED cells, blocking estrogen signaling led to increased tamoxifen resistance. Importantly, in all these cell lines (TAMR and LTED), methylation mediated downregulation of *ASS1* suggests role of estrogen signaling in protecting *ASS1* promoter methylation. Importantly, *ASS1* is not methylated and silenced in SUM44 parental cells as shown in our studies, suggesting a different mechanism of *de novo* resistance to anti-estrogen in this patient.

ASS1, the rate-limiting enzyme for the biosynthesis of arginine catalyzes the conversion of L-citrulline and aspartate to Arginosuccinate [40]. Downregulation of *ASS1* results in availability of aspartate for purine and pyrimidine biosynthesis facilitating cell proliferation (**Fig. 6E**). Zhou *et.al.* used Spinosyn A and its derivative LM-21 to augment ASS1 enzymatic activity that led to inhibition of cancer cell by blocking pyrimidine biosynthesis [41]. Our study showed enrichment of purine metabolism in the TAMR cells and in the *ASS1* knockdown cells suggesting that ASS1 could be both a biomarker as well as therapeutic target in tamoxifen resistant ILC. Use of decitabine, a demethylating agent augment *ASS1* expression and enhanced tamoxifen efficacy in our study. Decitabine is clinically approved to treat myelodysplastic syndrome [42], but demonstrated minimal efficacy as monotherapy in solid tumors [43, 44, 45]. However, combination therapy with targeted agents and chemotherapeutic agents have shown some promise [46], including a recently completed window of opportunity study [47]. Further studies are needed to see if prior treatment with decitabine can overcome tamoxifen resistance in ILC patients.

Association of low *ASS1* expression with poor overall survival has been reported in multiple cancer including bladder [48], myxofibrosarcoma [49] and breast cancer [50], although the number of breast cancer patients included were limited (n=149) and all subtypes were analyzed. However, analysis of 1980 breast cancer patients from METABRIC data set in our study revealed significant correlation of high *ASS1* with poor overall survival. Importantly, subtype specific analyses focusing on ILC have not been reported before. Correlation of low *ASS1* expression with poor OS in ILC patients only if treated with endocrine therapy as revealed in our study, highlights the potential of ASS1 as a biomarker for endocrine resistance in ILC. We cannot disregard the possibility that such correlation also exists for endocrine resistant IDC patients. Further studies are warranted to establish this relationship in IDC but beyond the scope of this study. Importantly, *ASS1* loss has also been implicated in chemotherapeutic resistance in different tumors including non-small cell lung cancer, ovarian cancer and hepatocellular carcinoma [49, 51, 52, 53, 54, 55, 56], suggesting ASS1 to be a vulnerable metabolic hub for development of therapy resistance.

Our study shows elevated phosphorylation of CAD, a key enzyme in pyrimidine biosynthesis, in both TAMR and *shASS1* cells. This further reinforces the notion that metabolic rewiring of existing pathway is a mechanism that cancer cells use to develop treatment resistance. Similar metabolic alteration was reported by Rabinovich et al. showing reduced ASS1 activity in cancer facilitates pyrimidine synthesis by activating CAD [57]. Our transcriptomics data further revealed heightened expression of several key genes involved in nucleotide biosynthesis pathways, suggesting that acquisition of drug resistance is a multiprong adjustment by the cancer cells for maximum benefit under the adversity of drug treatment. It is therefore likely that we could improve the efficacy of tamoxifen in TAMR cells by targeting nucleotide biosynthesis. As a proof of concept we used Farudodstat, to inhibit DHODH and observed synergistic effect when combined with tamoxifen. This combination therapy could be a potential treatment option for TAM-resistant ILC, warranting further investigation in the clinical setting.

Some limitations of our study warrant acknowledgment. First, our investigation is constrained by the availability of only two ILC cell lines, however, as noted above these are globally accepted as robust ILC models. Correlation of our finding with patient data partly addresses this limitation. The observed variability in experimental outcomes between these two cell lines may stem from their origin as discussed previously. In addition, metabolomic and transcriptomic studies provide a snapshot of the metabolic and gene expression state of the cells at the time of harvest. Although seeded at the same density, inherent difference in rate of cell growth between the lines in this study poses challenge in capturing the identical metabolic state for all the lines. This is reflected in the dissimilarities in pathway enrichment and metabolic intermediate levels observed in different cell pairs. However, the three pathways and *ASS1* alteration, highlighted in this study are common among the two ILC cell lines and could be a potential predictive marker in ILC patients.

In conclusion, our study highlights ASS1 as a potential biomarker for tamoxifen response and overall survival in ILC patients treated with endocrine therapy. Methylation mediated downregulation of *ASS1* provides an opportunity for clinical intervention of endocrine resistant ILC patients with demethylating agents, such as Decitabine and need to be explored further. Additionally, therapeutic interventions targeting nucleotide biosynthesis pathways show promise in overcoming tamoxifen resistance. Upregulation of purine and pyrimidine biosynthesis pathway enzymes in TAMR cells underscores the importance of metabolic adaptations in resistance. Further research into role of estrogen in protecting *ASS1* promoter methylation and therapeutic implications is expected to enhance treatment strategies of this understudied subtype of breast cancer.

## Materials and Methods

### Cell lines

ILC cell lines MDA-MB-134-VI (MB134) (ATCC, USA) and SUM44PE (SUM44) (Asterand, USA) were used to develop the tamoxifen-resistant (TAMR) cells by continuously exposing them to 100 to 500 nM of 4-hydroxy tamoxifen (4-OHT) for over 6 months. Parental and TAMR MB134 cells were grown in a 1:1 ratio of Dulbecco’s modified Eagle’s medium (DMEM; Gibco, USA) and Leibovitz’s L-15 medium (Gibco, USA), supplemented with 10% fetal bovine serum (FBS) and penicillin-streptomycin. Parental and TAMR SUM44 cell lines were cultured in Ham’s F-12 (Gibco, USA) supplemented with 1 g/L bovine serum albumin, 5 mM ethanolamine, 10 mM HEPES, 1 µg/mL hydrocortisone, 5 µg/mL insulin, 50 nM sodium selenite, 5 µg/ mL apo-transferrin and 10 nM triodo-L-thyronine. MB134TAMR and SUM44TAMR cells were maintained in media containing 100 nM and 500 nM 4-OHT respectively. MB134-LTED (Long Term Estrogen Deprived) and SUM44-LTED cells are generous gifts from Dr. Steffi Oesterreich (University of Pittsburgh) routinely maintained in IMEM + 10 % CS-FBS (Charcoal-Stripped FBS) and penicillin-streptomycin [34]. All cell lines were maintained at 37°C in a humidified 5% CO_2_ incubator. All cell lines tested Mycoplasma-free before the experiments.

### RNA isolation and sequencing

Total RNA was extracted from exponentially growing cells using Trizol (Invitrogen, USA) following manufacturer’s protocol. The TAMR cells were grown in tamoxifen-free media for 72 hours before harvest. RNA from three biological replicates of each cell line was subjected to RNA-seq analysis (Novogene, USA). Data processing and pathway analysis was performed by Novogene, and KEGG (Kyoto Encyclopedia of Genes and Genomes) pathway database, GO (Gene Ontology) database, and Reactome Pathway database were used for data analysis.

### Polar Metabolite Extraction

Exponentially growing cells (∼1 × 10^6) in quadruplet were used for metabolite extraction. Polar metabolites extraction was performed via a cold methanol extraction as previously described [58, 59]. Briefly, the cells were washed with cold phosphate buffered saline (PBS) followed by addition of 250 μL of methanol (LCMS-grade). Internal standards containing ^13^C and ^15^N labeled amino acids mix (1.2 mg/mL) were introduced to the samples in a volume of 50 μL. The cells were homogenized for 2 minutes and incubated at −20 °C for 20 minutes. The resulting homogenate was centrifuged to pellet the debris, and 150 μL of the supernatant was transferred to an LC-MS vial for further analysis. Additionally, a pooled quality control (QC) sample was created by combining an equal volume of all supernatants into a separate vial and mixed thoroughly using a vortex.

### LC-MS/MS System

Untargeted metabolomics was performed to uncover the metabolic alterations responsible for the drug-resistant phenotype using our established workflow [59]. The LC–MS/MS analyses were performed on a Vanquish ultra high-performance liquid chromatography (UHPLC) system (Thermo Scientific, Waltham MA, USA) coupled to a Qexactive™ Hybrid Quadrupole-Orbitrap™ Mass Spectrometer (Thermo Scientific, Waltham MA, USA). A sample volume of 5 μL was injected onto an Xbridge BEH Amide XP Column, 130Å (150 mm × 2.1 mm ID, particle size 2.5 μm) (Waters Corporation, Milford, MA, USA). The column oven was maintained at 40 °C. Mobile phase A consisted of a mixture of 5 mM NH_4_Ac in Acetonitrile/H_2_O (10:90, v/v) containing 0.1% Acetic acid. Mobile phase B consisted of 5 mM NH_4_Ac in Acetonitrile//H2O (90:10, v/v) containing 0.1% Acetic acid. The mobile phases were delivered at a flow rate of 0.3 mL/min for a 20-minute run with the following stepwise gradient for solvent B: firstly 70%; 0-5 min 30%; 5-9 min 30%; 9-11 min 70%. A divert valve was used to direct the flow to waste during the final 5 minutes of the run. The Qexactive™ was equipped with an electrospray ionization source (ESI) that was operated in both negative and positive ion modes to encompass a broader range of metabolite detection. The ESI source setting and the compound dependent scan conditions were optimized for full scan MS mode and ranged between 150 and 2,000 m/z. The ion spray voltage was set at 4 kV with a capillary temperature of 320°C. Sheath gas rate was set to 10 arbitrary units. Scans of 1ms were performed at 35,000 units resolution. A QC sample followed by a blank injection was introduced after every 10 biological sample injections. The pooled samples were leveraged for the top 10 MS/MS analyses, employing dynamic exclusion to identify compounds during the analysis.

### Growth Kinetics

Exponentially growing cells (30,000/well) were seeded in triplicate in 24-well plates. Subsequently, at 0, 24-, 48-, 72-, and 96-hour post-seeding, cells were trypsinized and counted using a cell counter (LUNATM, L12001). Fold change in growth was calculated with cell number at 0-hour timepoint as 1.

### Generation of *ASS1* knockdown cells

For shRNA-mediated knockdown of *ASS1*, lentivirus coding for *ASS1* shRNA in pLKO.1 backbone vector were purchased from Sigma (TRCN0000045554, sequence: GCCTGAATTCTACAACCGGTT). Exponentially growing MB134 and SUM44 cells (300,000/well) were seeded in 6-well plates. Overnight cultures were infected with 5-10 μL of viral particles in fresh medium containing 10 μg/mL polybrene. The cells were then incubated overnight followed by replacement of the virus containing media with fresh complete medium after 16 hours of incubation. Cells were expanded and transduced cells were selected using puromycin. The efficiency of viral infection in SUM44PE cells was assessed for GFP positivity using the EVOS M7000 imaging system. Knockdown of *ASS1* was validated by qRT-PCR and Western blot analysis. Similar experiments were conducted with the empty vector pLKO.1 to generate the control cells.

### Quantitative RT-PCR analysis

DNase treated total RNA was used to synthesize cDNA using High-Capacity cDNA Reverse Transcription Kit (Applied Biosystems). qRT-PCR was performed in triplicate using 96-well StepOne Real-Time PCR System. 36B4 was used a housekeeping gene. Primer sequences are: ASS1-F: GCTGAAGGAACAAGGCTATGACG and ASS1-R: GCCAGATGAACTCCTCCACAAAC. 36B4-F: GGTTGTAGATGCTGCCATTGTC and 36B4-R: GCCCGAGAAGACCTCCTTTTTC.

### Western blot analysis

Whole cell lysates in radioimmunoprecipitation assay (RIPA) buffer (50 mM Tris-HCl pH 7.4, 150 mM NaCl, 1% NP-40, 0.1% SDS; Sigma-Aldrich), supplemented with protease and phosphatase inhibitors (Sigma) were resolved on SDS polyacrylamide gels. Following electrophoresis, proteins were transferred onto 0.45 µM PVDF membranes. Nonspecific binding was blocked by incubation with blocking buffer (Rockland) for 60 min at room temperature. The membranes were probed for ASS1 [Cell Signaling Technology (CST), 70720], GAPDH (CST, 2118S), ERα (Abcam, ab32063), HER2 (CST, 2242) and Phosphor-CAD (Ser1859) (CST,1266). ASS1 and GAPDH was detected using IR800CW dye-tagged IgG secondary antibody (LICOR, 926-32211). Phosphorylated proteins (p-CAD) were detected using peroxidase-conjugated anti-rabbit secondary antibody (CST, 7074) and enhanced chemiluminescence western blot detection reagents (Pierce, Thermo Scientific). The Odyssey CLx and Fc systems (LI-COR Biosciences, USA) were used for western blot imaging. All original western blot images are provided in **Supplementary file 2.**

### Methylation-specific PCR

Computational analysis using ‘CpG Island Finder’ (CpG Islands (bioinformatics.org) was performed to locate CpG island on gene promoter. DNA (200–500 ng) extracted from exponentially growing cells were subjected to bisulfite conversion using the EZ DNA methylation kit (Zymo Research Corporation, USA). Treatment of genomic DNA with sodium bisulfite converts unmethylated cytosine residues to uracil, while methylated cytosine residues remain unchanged. The methylation status of *ASS1* promoter was determined using methylation-specific PCR (MSPCR). Primers for MSPCR were designed using MethPrimer software (https://www.urogene.org/methprimer/). The bisulfite-converted DNA (1-4 µL) was used for PCR reactions with primers specific for either methylated (F: GTCGGTATCGGATAGAAGTGAGTAC, R: ATAACTCAAAAACGAAAAATAACCG) or unmethylated sequences (F: TTGGTATTGGATAGAAGTGAGTATGA, R: AACTCAAAAACAAAAAATAACCACA)[54]. PCR conditions were as follows: 8 cycles of 95 °C for 2 min, 61 °C for 30 s, and 72 °C for 30 s, followed by 32 cycles of 95 °C for 30 s, 61 °C for 30 s, and 72 °C for 30 s, with a final extension at 72 °C for 5 min. PCR products were electrophoresed in 2% agarose gels and visualized using a transilluminator.

### Cell viability assay

MTT assay kit (Roche) was used to assess cell viability and drug effect. Fifteen thousand cells were seeded per well in triplicates in a 96 well plate, and overnight cultures were treated with drugs for 120 hours. This was followed by addition of MTT reagent and solubilizing agent following manufacturer’s protocol. The drugs included 5 µM of 5-Aza-2 deoxycytidine (dAzaC, Sigma), also known as Decitabine, 4-hydoxy tamoxifen (0 – 25 µM, Cayman chemical) and Farudodstat (7.5 – 20 µM, Cayman Chemical), a pyrimidine biosynthesis inhibitor. CompuSyn 1.0 (https://compusyn.software.informer.com/) was used to analyze combinatorial effect of two drugs. Combination index (CI) value = 1, <1 and >1 indicate additive, synergistic and antagonistic effect, respectively.

### Migration assays

*In-vitro* cell migration assays were conducted using Transwell chambers (Corning, USA) coated with collagen (50μg/mL) on the exterior of the inserts for 60min at 37°C. For the migration assay, 500,000 cells in serum free media were seeded in the inserts. A chemotactic gradient was established by adding 0.6 mL of complete growth medium containing 10% FBS in the lower chamber. After 120 hours of incubation the unmigrated cells in the inserts were removed using a cotton swab, the migrated cells were fixed in methanol, and stained with 0.5% crystal violet solution. The area occupied by migrated cells was quantified using ImageJ software.

### Patient dataset analysis

Human breast cancer patient data was obtained from The Cancer Genome Atlas Firehose BRCA cohort (TCGA_BRCA) and the METABRIC invasive breast carcinoma cohort were obtained from the cBioPortal webpage. Analysis of METABRIC patient data was conducted in R 4.2.2 utilizing dplyr 1.0.10, tibble 3.1.8, ggplot2 3.4.0, and survival 3.5.8 packages to import, subset, and analyze relevant patient data[60]. The Kmplotter webtool was used to analyze the Kmplotter meta cohort (Kmplotter.com) [61].

### Statistical analysis

Three independent replicates of all the experiments were conducted. Western blot experiments were independently repeated using cell lysates from three biological replicates and data expressed as the mean ± standard deviation (SD). Statistical analyses between two groups were performed using Student’s t-test. One-way ANOVA was used for multiple group comparisons. Percentage of cell viability and IC_50_ value were calculated using GraphPad Prism. Differences in survival of patients from the METABRIC cohort and the KM plotter meta cohort were determined via log-rank test. A *p*-value < 0.05 was considered statistically significant.

#### Metabolomic Data Processing and Statistical Analysis

Initial screening of the spectral peaks was performed using the Quan browser module of Xcalibur version 4.0 (Thermo Fisher Scientific, Waltham, MA, USA). The MS data were searched against our in-house database containing experimentally obtained MS/MS spectra of 171 authentic analytical standards using Compound Discoverer software (Thermo Scientific, San Jose, CA, USA). The raw data was normalized to the protein content per replicate. Subsequently, the spectra underwent filtration to diminish redundancy and ensure instrument reproducibility. Any metabolite exhibiting a coefficient of variation exceeding 20% was eliminated before subsequent analysis. Statistical analyses, including univariate T-test, were conducted using the online resource MetaboAnalyst 5.0. Partial Least Square Discriminant Analysis (PLS-DA) was employed to interpret the metabolic variances between the sensitive and resistant cell lines. VIP (Variables Important in Projection) plots were generated to visualize the key metabolites contributing to the deregulated metabolic processes. Overall metabolite data was subjected to quantitative enrichment analysis to pinpoint the deregulated metabolic process within the cell pairs. A Venn diagram was generated to show the distinguishable gene expression profiles among samples and summarize the mutually deregulated pathways across both metabolomics and transcriptomics datasets. The gene-metabolite interaction networks were constructed by integrating annotated metabolites or genes with comprehensive interaction data from Search Tool for Interactions of Chemicals (STITCH) [62]. This tool utilizes interaction data from peer-reviewed literature to assess node importance within the network based on degree centrality and betweenness centrality. Degree centrality is determined by the number of connections a node has with others, while betweenness centrality calculates the number of direct routes passing through the node. These measures help identify metabolic hubs within the network.

## Acknowledgements

We are grateful for the philanthropic support of Helen and Brad Anderson of Columbus, Ohio towards this research and grateful to Steffi Oesterreich, Ph. D of University of Pittsburgh for providing the ILC-LTED cell lines.

## Author contribution statement

AG and FC: Experimental design, Methodology, Investigation and Writing Original Draft. JR: Bioinformatic analysis of patient data, Writing and Review. NP, AK, ES: Investigation. STS: Supervision of bioinformatics analysis and Review. JZ: Resources, Data curation, Editing. DGS: Supervision and Editing. BR: Funding acquisition, supervision, editing and project administration. SM: Conceptualized, Supervision, Data interpretation, Writing-Review and Editing. All authors read and approved the final manuscript.

## Ethics Statement

This study was conducted using human invasive lobular carcinoma (ILC) cell lines. All cell lines used were obtained from accredited sources (ATCC) and handled according to standard laboratory safety and ethical guidelines.

## Availability of Data and Materials

Both the RNA sequencing and Metabolomics data generated in this study will be submitted to the publicly available databases [Gene Expression Omnibus (GEO) database-for RNA seq data and https://massive.ucsd.edu/ProteoSAFe/static/massive.jsp -Metabolomics data] upon acceptance of the manuscript and before final printing. The data supporting the findings reported in this study are available from the corresponding author upon reasonable request.

