## Supplementary Figures and tables for "Invasive lobular carcinoma integrated multi-omics analysis reveals silencing of Arginosuccinate synthase and upregulation of nucleotide biosynthesis in tamoxifen resistance"

### Supplementary Fig. S1

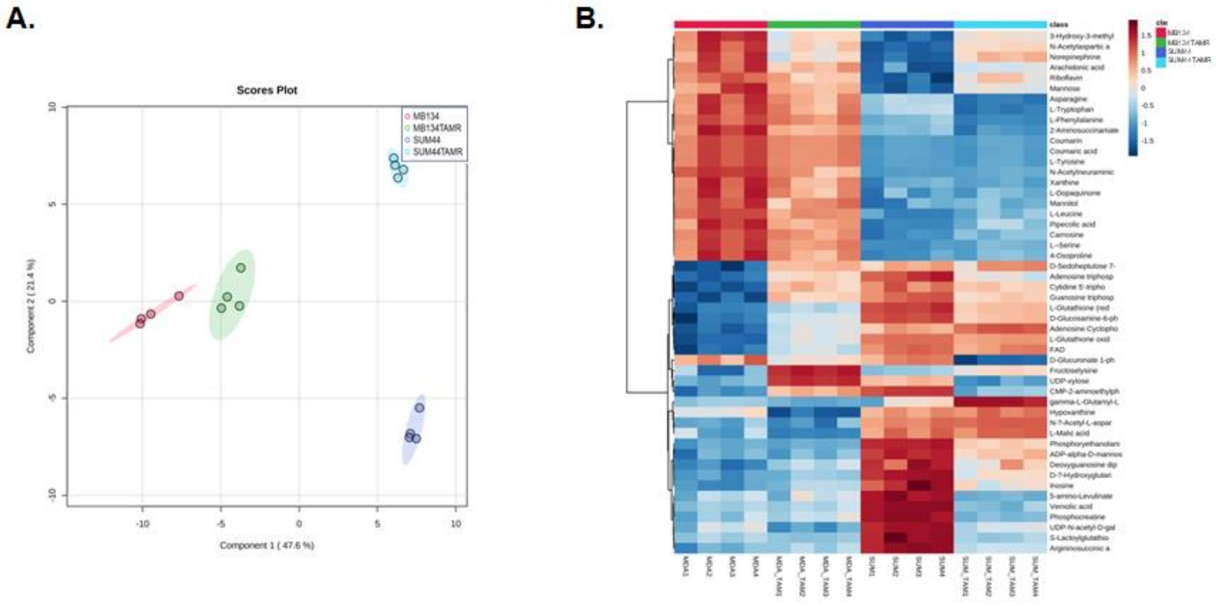


**Supplementary Fig. S1**: Mutually deregulated metabolic pathways between parental and TAMR ILC cell lines. **A.** Partial Least Squares Discriminant Analysis (PLS-DA) comparing the overall metabolic profiles of the parental and TAMR ILC cell lines, illustrating the separation of metabolic profiles between the four cell lines. **B.** Heatmap depicting the relative abundance of each metabolite across the parental and TAMR ILC cell lines. Each row represents a metabolite, while each column represents a cell line. The color intensity corresponds to the abundance level of each metabolite, with red indicating higher abundance and blue indicating lower abundance (scale shown). The heatmap highlights the alterations in metabolic pathways that are deregulated between the parental and tamoxifen resistant ILC cell lines.

### Supplementary Fig. S2

A.

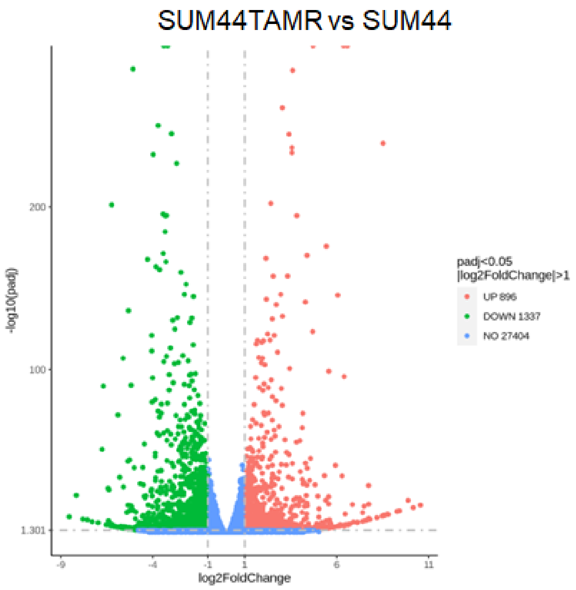


B.

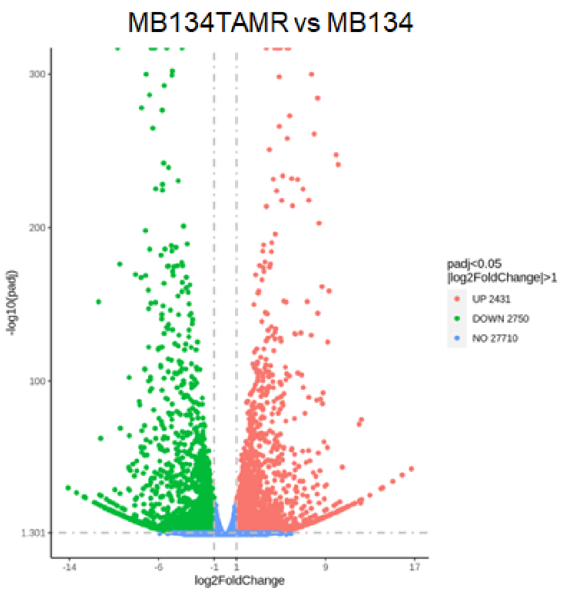


**Supplementary Fig S2:** RNA seq analysis. **A.** Volcano plot of RNA sequencing data showing differential expression of genes and their statistical significance in SUM44 vs. SUM44TAMR, and **B.** MB134 vs. MB134TAMR cell lines.

### Supplementary Fig. S3

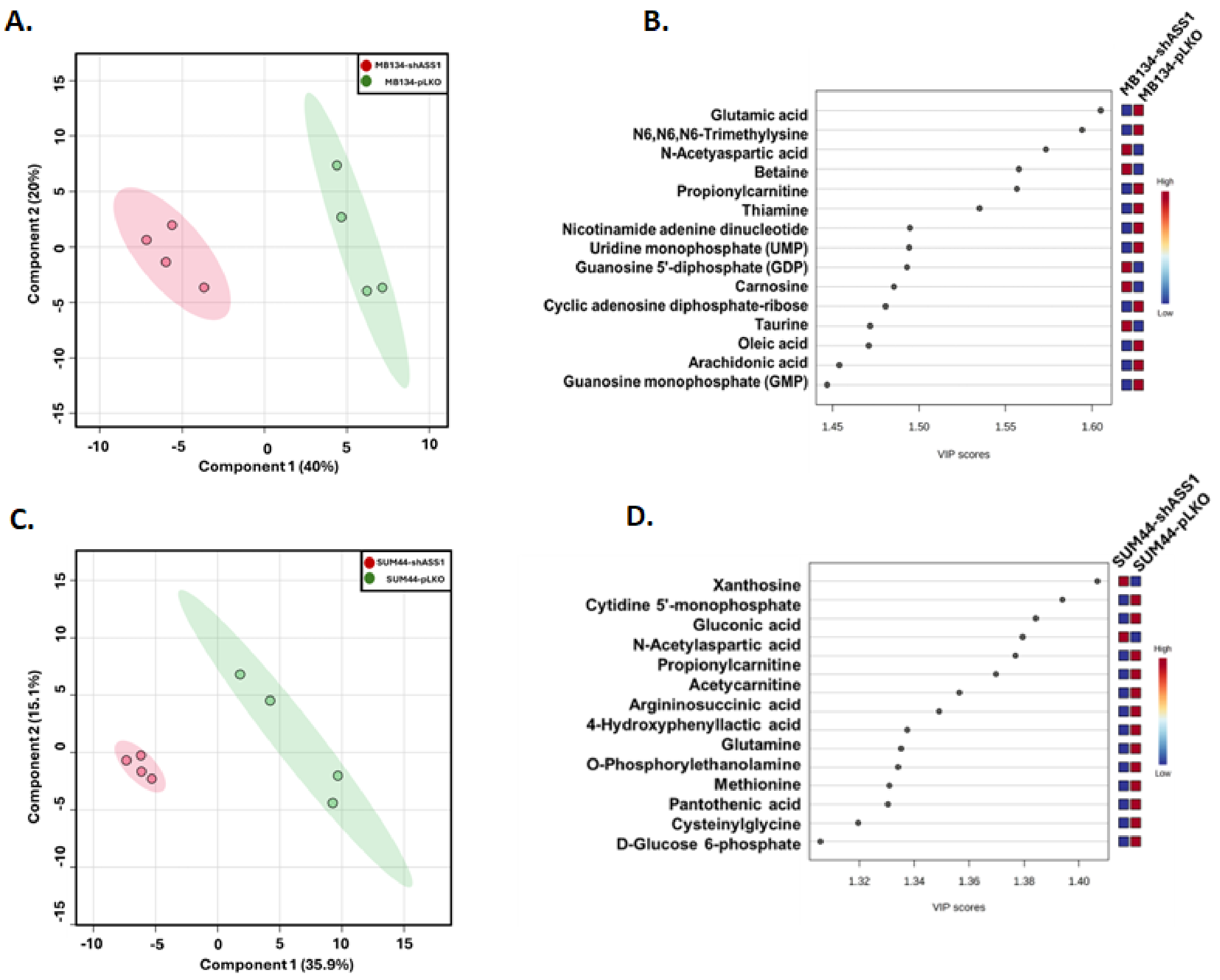


**Supplementary Fig S3: A.** Partial Least Squares Discriminant Analysis (PLS-DA) plot comparing metabolic profile of MB134-*shASS1* vs. MB134-pLKO cells. **B.** Variance Importance in Projection (VIP) plot highlighting the top metabolites driving the separation of the MB134-*shASS1* vs. MB134-pLKO cells. **C.** PLS-DA plot comparing metabolic profile of SUM44-*shASS1* vs. SUM44-pLKO cells. **D.** VIP plot highlighting the top metabolites driving the separation of the SUM44-*shASS1* vs. SUM44-pLKO cells.

#### Supplementary Fig. S4

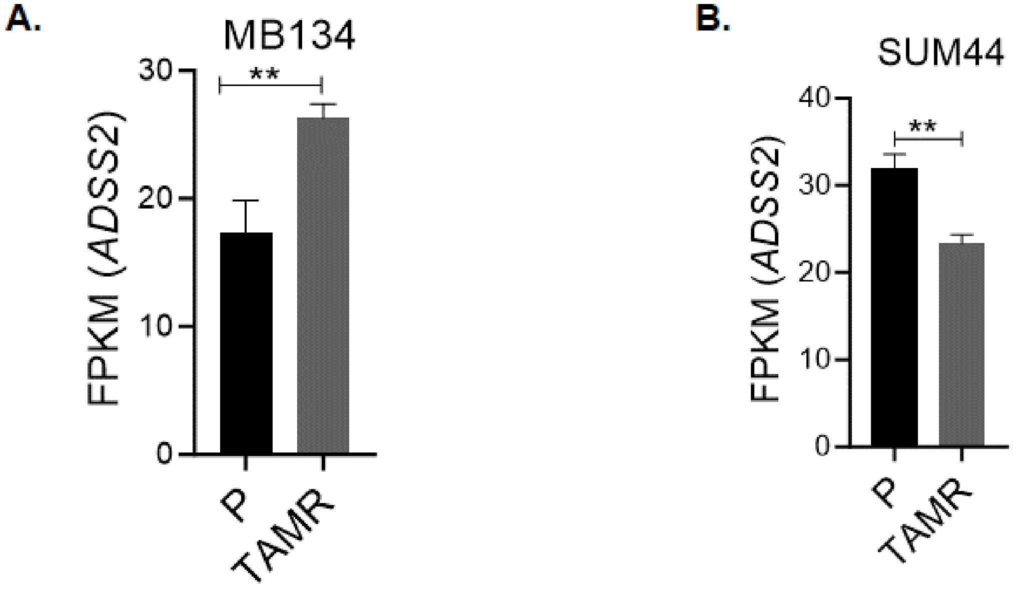


**Supplementary Fig S4. A.** Expression analysis of *ADSS2* in MB134TAMR vs. MB134 and **B.** SUM44TAMR vs. SUM44 cell lines using Fragments Per Kilobase of transcript per Million mapped reads (FPKM) from RNA sequencing data. P- Parental, TAMR-Tamoxifen resistant

**Supplemental Table 1:** A full list of the significantly deregulated pathways between the queried cell pairs.

| Pathway Name | -log(p) | Impact | Cell pair |
|---|---|---|---|
| Alanine, aspartate and glutamate metabolism | 7.57 | 0.47 | SUM44PE |
| D-Glutamine and D-glutamate metabolism | 6.02 | 0.50 | SUM44PE |
| Arginine and proline metabolism | 4.78 | 0.33 | SUM44PE |
| Arginine biosynthesis | 4.33 | 0.37 | SUM44PE |
| Lysine degradation | 4.30 | 0.28 | SUM44PE |
| Purine metabolism | 4.04 | 0.28 | SUM44PE |
| Linoleic acid metabolism | 3.92 | 1.00 | SUM44PE |
| Vitamin B6 metabolism | 3.74 | 0.49 | SUM44PE |
| Glycine, serine and threonine metabolism | 3.66 | 0.22 | SUM44PE |
| Tyrosine metabolism | 3.64 | 0.36 | SUM44PE |
| Ascorbate and aldarate metabolism | 3.43 | 0.50 | SUM44PE |
| Pentose and glucuronate interconversions | 3.43 | 0.20 | SUM44PE |
| Arachidonic acid metabolism | 2.95 | 0.31 | SUM44PE |
| Riboflavin metabolism | 2.66 | 0.50 | SUM44PE |
| Alanine, aspartate and glutamate metabolism | 6.24 | 0.47 | MDA-MB134 |
| Purine metabolism | 5.16 | 0.28 | MDA-MB134 |
| D-Glutamine and D-glutamate metabolism | 4.93 | 0.50 | MDA-MB134 |
| Pentose phosphate pathway | 4.44 | 0.25 | MDA-MB134 |
| Lysine degradation | 3.92 | 0.28 | MDA-MB134 |
| Arginine biosynthesis | 3.88 | 0.37 | MDA-MB134 |
| Riboflavin metabolism | 3.88 | 0.50 | MDA-MB134 |
| Tyrosine metabolism | 3.68 | 0.36 | MDA-MB134 |
| Arginine and proline metabolism | 3.23 | 0.33 | MDA-MB134 |
| Arachidonic acid metabolism | 2.48 | 0.31 | MDA-MB134 |
| Phenylalanine, tyrosine and tryptophan biosynthesis | 2.25 | 1.00 | MDA-MB134 |
| Phenylalanine metabolism | 2.25 | 0.36 | MDA-MB134 |
| Glycine, serine and threonine metabolism | 2.02 | 0.22 | MDA-MB134 |

**Supplementary Table 2:** List of genes involved in the three dysregulated pathways

| Pathways | Genes |
|---|---|
| **Alanine_Asparate_Glutamine & Purine** | ADSS2 |
| | ADSL |
| | ADSS1 |
| | PPAT |
| **Alanine_Asparate_Glutamine & Arginine_Proline** | GOT2 |
| | ALDH4A1 |
| | GLS |
| | GLUD1 |
| | GLUL |
| | ASS1 |
| | GOT1 |
| | ASL |
| | CPS1 |
| | GLUD2 |
| | GLS2 |
